# Constrained flexibility of parental cooperation limits evolutionary responses to harsh conditions

**DOI:** 10.1101/2021.04.02.438190

**Authors:** J.B. Moss, A.J. Moore

## Abstract

Parental care is predicted to evolve to mitigate harsh environments, thus adaptive plasticity of care may be an important response to climate change. In biparental species, fitness costs may be reduced with plasticity of behavior among partners. We investigated this prediction with the burying beetle, *Nicrophorus orbicollis*, by exposing them to contrasting benign and harsh thermal environments. We found strong fitness costs under the harsh environment, but rather than select for more care, visualized selection was stabilizing. Examining different components of care revealed positive directional selection gradients for direct care and strong stabilizing selection gradients for indirect care, resulting in constrained evolutionary responses. Further, because males and females did not coordinate their investments, the potential for adaptive plasticity was not enhanced under biparental care. Females cared at capacity with or without male partners, while males with partners reduced direct care but maintained indirect care levels. Decision rules were not altered in different environments, suggesting no shift from sexual conflict to cooperation. We suggest that the potential for parenting to ameliorate the effects of our climate crisis may depend on the sex-specific evolutionary drivers of parental care, and that this may be best reflected in components of care.

## Introduction

Parental care is expected to evolve to mitigate hostile and unpredictable environments (Wilson 1975). However, the extent that ecological conditions further modify parenting once it evolves may depend on plasticity of parental care in response to environmental stress; that is, the phenotypic variation exposed to selection. One potential source of plasticity of care where such variation might be exposed is biparental cooperation. Theoretically, the default of biparental systems is sexual conflict over which parent cares (Lessells 2012), which can lead to overall care deficits (McNamara et al. 2003, Lessells and McNamara 2012) and, ultimately, to one parent being as effective or more effective at caring for offspring than two parents (Clutton-Brock 1991, Smiseth et al. 2005, Trumbo 2006). However, the joint rearing of offspring may also allow parents to breed under harsh conditions that would otherwise constrain single parent breeding (Wilson 1975, Emlen 1982). This is because: 1) with more than one caregiver there is more scope for increasing total care allocation (i.e., additive care; Ratnieks 1996, Clutton-Brock et al. 2001, Johnstone 2011, Savage et al. 2013), and 2) the efforts of a second parent may offset some costs of care to the primary caregiver (i.e., load lightening; Crick 1992, Johnstone 2011). If true, then transitions to stable biparental care and an increased capacity for cooperation should coincide with expansion into increasingly harsh environments (Wesolowski 1994, 2004).

To date, tests of the ‘hostile environment’ hypothesis as it relates to cooperation over offspring rearing have produced equivocal results (Wynne-Edwards and Timonin 2007, AlRashidi et al. 2010, 2011, Öberg et al. 2015, Remeš et al. 2015, Wiley and Ridley 2016, Shen et al. 2017, Vincze et al. 2017, Guindre-Parker and Rubenstein 2018, Lejeune et al. 2019, Lin et al. 2019, Vági et al. 2020). However, the vast majority of insights derive from studies of birds – a group for which biparental care is nearly ubiquitous and rarely decoupled from social monogamy (Cockburn 2006). Hence, while reports of enhanced pair coordination under adverse conditions (i.e., approaching egalitarian division of labor) accumulate from the bird literature (AlRashidi et al. 2010, 2011, Vincze et al. 2017), the extent to which such causal links are generalizable across taxa is unclear. Yet transitions to biparental care have occurred repeatedly outside of the avian tree, including in diverse vertebrate (Reynolds et al. 2002) and invertebrate lineages (Trumbo 2012, Suzuki 2013, Gilbert and Manica 2015). Such systems offer rich opportunities to expand the taxonomic scope of investigations into the factors that shape biparental care dynamics.

Burying beetles (Genus: *Nicrophorus*) provide an ideal complement to avian systems for investigating the mechanisms of cooperation and conflict over offspring care (Smiseth 2019), particularly in the context of environmental stress and plasticity. First, burying beetle parental care reflects their ecology. The beetles breed on an ephemeral and widely desirable resource, a dead vertebrate, resulting in both rapid development and parental care involving direct provisioning of predigested food and defense of the developing young (Eggert and Müller 1997, Scott 1998a). Burying beetles are also subsocial; they do not form social associations outside of brief periods of parental care. Therefore, unlike most vertebrates, sources of variation in parental investment can be readily dissociated from other pervasive aspects of social life. Second, we know that there is capacity for plasticity of species that show biparental care because burying beetle males do not work at their maximum. Indeed, parental care of burying beetles is sex-biased, with females performing the majority of total caregiving duties (Eggert and Müller 1997, Smiseth and Moore 2004, Benowitz and Moore 2016) while males provide less direct care in the presence of a female partner (Parker et al. 2015, Pilakouta et al. 2018). The quantity or composition of female behavior does not depend on the presence of a male. However, males of this genus are also highly flexible and capable of adopting larger parental roles as needed to compensate for compromised partner state (e.g., partner loss (Trumbo 1991, Smiseth et al. 2005, Suzuki and Nagano 2009, Parker et al. 2015, Cunningham et al. 2019), handicapping (Creighton et al. 2015), or inbreeding level (Mattey and Smiseth 2015). Finally, many burying beetles are flexible in the social form of parenting they provide, with uniparental female care, uniparental male care, and biparental care all expressed within natural populations (Trumbo 1991, Scott 1998a, Smiseth and Moore 2004, Suzuki and Nagano 2009, Benowitz et al. 2016). If it is true that multiple parents provide more effective care to offspring in hostile conditions (i.e., through additive and/or load-lightening effects), then members of the more flexible sex should also be less inclined to withhold care in response to a generalized environmental stressor, which may compromise the states of both parents.

Here, we use *Nicrophorus orbicollis* – a primarily biparental species and among the few members of the temperate species complex to have successfully expanded into the warmer climate of the US southeast (Trumbo 1990) – to examine the role that plasticity of parental investment plays in mitigating harsh ambient conditions. High temperatures, as occur at low latitudes, are generally implicated in more costly and less profitable reproduction in burying beetles (Meierhofer et al. 1999, Müller et al. 2007, Steiger et al. 2007, Jacques et al. 2009, Quinby 2016, Ong 2019, Feldman 2020). Individuals breeding under these conditions have been found to suffer reduced lifespans and lower lifetime reproductive success (Laidlaw 2015). We used a mixed factorial design with repeated measures to test whether beetles acclimated to high temperature (i.e., harsh) breeding conditions respond by increasing parental investment. Further, we quantified within-subject behavioral comparisons to examine the extent that sex-specific plasticity and the capacity for biparental care drives responses to hostile environments. Contrary to our prediction that plasticity of care will allow *N. orbicollis* to tolerate higher temperatures, we find that there is stabilizing selection on total care with an optimum independent of the form of care provided. This occurs because the components of care are under different forms of selection, the components are not independent, and individual variation did not reflect a plastic response to subtle variation in their partner’s behavior.

## Methods

### Study System

*Nicrophorus orbicollis* is a large-bodied, ecological generalist that breeds on small (~20 g) to medium (~100 g) vertebrate carcasses in North American woodlands. The species has a large latitudinal distribution (from southern Canada to northern Texas), with breeding seasons at the southern margins characterized by higher temperatures (3–8°C on average) and a greater frequency of reproductive failure (Trumbo 1990). As with most members of the genus, parental care is elaborate and extends into the post-hatching stage (Eggert and Müller 1997, Scott 1998a). During pre-hatching stages, parents work together to clean and prepare the carcass by removing hair and applying anal secretions to prevent microbial growth. During the post-hatching stage, parents continue to maintain the brood ball and also directly provision to begging young *via* regurgitation. While larvae of most burying beetles can survive without parents (Schrader et al. 2015, Jarrett et al. 2018), *N. orbicollis* show obligate parental care, meaning that larvae depend on direct provisioning for survival (Trumbo 1992, Capodeanu-Nägler et al. 2016, 2018). Parental care is described as predominantly biparental on the basis that males and females typically overlap with each other in the post-hatching stage (in 66% of cases; Benowitz and Moore 2016), and both sexes perform the full repertoire of parenting behaviors (Scott and Traniello 1990, Trumbo 1991, Scott 1998a). However, as is the case with any reproductive systems studied in detail, individual investment is highly flexible and subject to environmental and social pressures (Trumbo 1991, Scott 1998b, Creighton et al. 2015).

### Field collection and husbandry

*Nicrophorus orbicollis* were captured from Whitehall Forest, Athens GA, in the summer of 2020. Beetles were baited into hanging traps with salmon and collected twice weekly to breed an outbred laboratory colony. Simultaneously, Thermochron® iButton temperature loggers (©Maxim Integrated Products, Inc., San Jose, CA, U.S.A) were deployed ~10–12 cm underground at trap locations throughout our collection site to estimate the range of temperatures beetles likely experience in their subterranean brood chambers. *Nicrophorus orbicollis* begin emerging from hibernation in early spring and reach peak densities around midsummer (between late June and early August; Ulyshen and Hanula 2004). In 2020, mean daily temperatures during these two potential breeding windows – late spring/early summer (31 May–03 July) and mid/late summer (15 July–22 August) – ranged between 21.71±1.38°C and 23.82±0.77°C, respectively (Fig S1). Diurnal temperature fluctuations were between 0.75°C and 7.71°C. To capture this variation in the laboratory, we programmed two incubators to ramp between set points of diurnal temperatures over the course of 10:14 hour reverse light: dark cycles, simulating early and late summer breeding conditions, respectively. The first treatment, hereafter the ‘benign’ thermal environment, was set to ramp between 19°C (night) and 20°C (day), while the second, hereafter the ‘harsh’ thermal environment, was set to ramp between 23°C (night) and 24°C (day). Focal individuals for the experiment were selected from F01 and F02 colony lines, which were bred on countertops at room temperature (20±0.5°C). Larvae were divided evenly between the treatment incubators on the third day of pupal development to facilitate acclimation (adults eclosed into the environment in which they would ultimately breed) while controlling for possible early developmental effects of temperature. All virgins selected for the experiment were at least 14 days of age.

### Breeding trials

Breeding trials were carried out between October 2020 and January 2021. We used a mixed factorial design as outlined in Figure 1, in which social condition (uniparental or biparental) was measured as a within-subject factor and thermal environment (Benign or Harsh) was measured as a between-subject factor. The goal was to achieve a balanced experimental design with respect to the number of individuals undergoing repeated trials (N = 20 males and females per thermal environment), which would allow us to explicitly quantify differences in individual plasticity between the two thermal environments. To facilitate this, we randomized the order in which focal individuals were exposed to either social condition. To create the biparental condition, individuals were paired to a focal individual of the opposite sex within the same thermal environment. To create the uniparental condition, individuals were paired to a random unrelated beetle of the opposite sex (also within the same thermal environment) who would be removed between egg laying and hatching. Individuals who successfully completed their first trial would continue on to a second trial in the opposite social condition, while individuals that failed their first trial within seven days of pairing were restarted. Beetles were allowed one failure on their first attempt.

**Figure 1:**
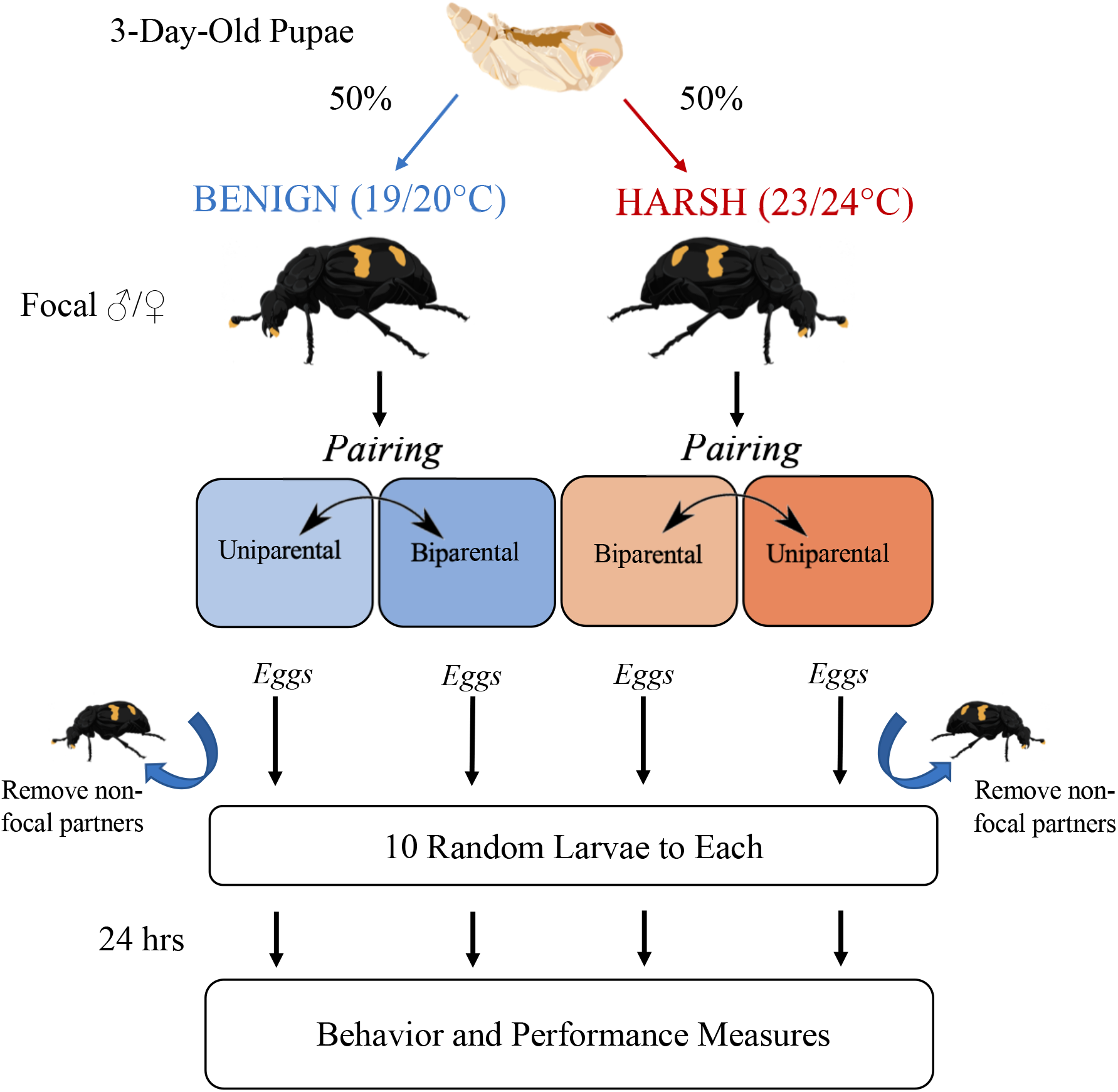
Schematic of mixed factorial experimental design. Focal individuals of each sex were divided evenly between the treatment incubators on the third day of pupal development to facilitate thermal acclimation (thermal environment = between-subject factor). At pairing, individuals were assigned randomly to a starting social condition (uniparental or biparental) and restarted in the opposite social condition upon successful completion of a first trial (social condition = within-subject factor). In uniparental trials, non-focal parents were removed after egg laying. All eggs were collected prior to hatching and each widowed parent or biparental pair was allocated a standardized number of larvae (N=10) of random genetic origin. Behavioral and performance measures were collected starting 24 hours into care.

At the start of each trial pairs were placed in a plastic box (17.2 x 12.7 x 6.4 cm; Pioneer Plastics, Dixon, KY, USA) filled with approximately two cm of moistened potting soil and containing a freshly thawed mouse carcass between 40 and 45 g (RodentPro, Evansville, IN, USA). Boxes were returned to the incubator where they were kept on a darkened shelf beneath blackout curtains to simulate an underground breeding environment. From pairing, breeding boxes were checked twice daily for eggs. Pairs with no eggs after seven days were restarted on a new mouse. Two days after eggs were first recorded, the brood ball and focal beetle(s) were transferred to a new breeding box (non-focal parents were removed) such that eggs could be collected and counted. This step was performed to facilitate brood standardization, which ensured that comparisons of performance would be attributed to parental care rather than differences in fertility or genetic quality. Eggs were placed in petri dishes with damp filter paper and monitored every 8 hours until larvae appeared. At this stage, synchronously hatching broods were randomly mixed, and each pair of fertile parents was given exactly 10 larvae. Broods that failed to hatch within five days of laying were recorded as unfertilized, and the pair was restarted.

### Behavior and performance measures

Behavioral observations were carried out 24 hours after introducing larvae, as previous work indicates that offspring provisioning peaks around this time (Smiseth et al. 2003). Breeding boxes were placed in a dark, temperature-controlled observation room (20°C) and allowed to acclimate for 30 minutes, ensuring that observed differences in parenting could not be attributed solely to temperature-dependent activity. Observations took place under red light over a 30-minute period. Behaviors were recorded every minute *via* instantaneous scan sampling. These included any instances of direct provisioning (i.e., mouth-to-mouth contact suggesting regurgitation of food to larvae), oral pre-treatment of feeding substrates (i.e., opening up the cavity or self-feeding to facilitate subsequent regurgitations), offspring association (i.e., in physical contact with larvae but not provisioning), carrion maintenance (i.e., positioned under brood ball or walking over brood ball exuding antimicrobial secretions), self-grooming, and off brood ball. Individual behaviors were then grouped into two broad categories – direct care (direct provisioning, oral pre-treatment of feeding substrates, and offspring association) and indirect care (carrion maintenance) – to arrive at 2x budget scores (ranging between 0 and 30) for each parent in each trial. “Self-grooming” and “off brood ball” were regarded as non-caring behaviors and were interpreted only in the inverse. For biparental pairs, total direct and indirect care were calculated by summing the time budget scores of the two parents.

After completing observations, brood boxes were returned to incubators and subsequently checked three times per day for parental desertion. Desertion was inferred when beetles were observed buried in the dirt away from the brood ball for three consecutive observations (Hopwood et al. 2015, Parker et al. 2015, Benowitz and Moore 2016). At this point, beetles were removed, and we recorded the duration of care (in days). Final weights were taken for each beetle at the end of a breeding trial, and those due for a second trial were fed and returned to the incubator for 1–2 days prior to restarting. To calculate and compare performance across trials, we measured two traits implicated in parental performance: total number of offspring surviving to the end of a breeding trial and mean larval mass (Parker et al. 2015). These measures were taken only after larvae dispersed naturally from the brood ball, to ensure maximal feeding time.

### Statistical analyses

All statistical analyses were conducted in R v. 4.0.3 (R Core Development Team 2019) using the package lme4 (Bates et al. 2015). We first examined evidence for fitness costs associated with the high temperature environment. Because a large number of deaths were recorded over the course of our experiment, our first analysis was of parental longevity. We used a Cox proportional hazard regression model implemented in the R package ‘survival’ (Therneau and Lumley 2015) to test the association between thermal environment and mortality, adjusting for sex. Our second analysis was of reproductive parameters. We used two-tailed *t*-tests to contrast the means of fecundity, fertility, and development time across all trials. To compare realized performance (number and mean mass of dispersing larvae), we used simple linear regression with thermal environment as a main effect and breeding history (binary specifying at least one previous breeding success between parents) as a covariate, to account for variation in parental experience within our study design.

After identifying costs associated with thermal stress, we split the dataset by thermal environment and examined evidence for variation in behaviors under selection. We explored how care influenced fitness through environment-specific performance gradients. We first plotted the relationship between standardized offspring mass and care allocation by social condition. Overall selection was visualized using total care (sum of all direct and indirect care). We next calculated standardized selection gradients (Lande and Arnold 1983, Brodie et al. 1995) for different components of care using number of larvae and mean larval mass to estimate fitness. We examined both linear and non-linear selection on components of parenting (maximum number of days parents attended, cumulative direct care, and cumulative indirect care). We then examined intra-environmental variation in parenting as a function of social condition by performing a mixed model analysis of variance (ANOVA) for each metric followed by specified *a priori* pairwise contrasts, comparing uniparental and biparental care behavior within a sex, implemented using the lsmeans package (Lenth 2016). To control for repeated measures of focal parents, we included male and female IDs as random block effects in the model design.

Our final analysis examined variation among individuals. We asked whether variation of parenting behavior reflected individual plasticity by examining patterns of individual behavior across the two thermal acclimation environments. To achieve this, we split the dataset by sex, retaining only individuals with repeated measures of behavior, and quantified how much behavioral variation *within* environments could be attributed to social plasticity. We first performed separate repeated-measures ANOVAs for each sex and thermal environment, using social condition as the within-subject factor and individual ID nested within trial number as the error term. To understand how individual male and female behaviors varied *between* environments, we carried out repeated-measures multivariate ANOVAs (RM MANOVA) using the R package, MANOVA.RM (Friedrich et al. 2019). The three behavioral metrics were specified as response variables and thermal environment was specified as a between-subject factor. To facilitate comparisons of duration of care given temperature-dependent larval development rates, attendance times were divided by the mean development times within each thermal environment. Wald-type statistics (WTS) and resampling *P-*values are reported for within- and between-subject factors and their interaction.

## Results

Over the course of this experiment, we initiated 358 breeding trials spread over two thermal environments and three social conditions. Only 165 trials resulted in larvae that survived through the 24-hr behavioral observation period. These included 32 biparental pairs, 29 uniparental males, and 30 uniparental females in the harsh environment, and 26 biparental pairs, 23 uniparental males, and 25 uniparental females in the benign environment. The final number of focal individuals with repeated measures amounted to 18 males and 20 females in the harsh environment and 20 males and 20 females in the benign environment.

### Fitness Costs of Thermal Stress

We observed strong adverse effects on fitness associated with thermal stress. Focal beetles in the harsh environment suffered a 47.6% higher mortality risk compared to counterparts in the benign environment (95% CI (1.43, 3.09), *P* < 0.001; Fig 2), with males outliving females (HR = 0.62, 95% CI (0.43, 0.90), *P* = 0.013). Within breeding trials, mortality accounted for 21.7% of failures among inexperienced breeders, and 31.6% among experienced breeders (compared to 12.9% and 10.2% in the benign environment, respectively). Reduced fecundity and infertility were other sources of breeding failure, with beetles laying fewer (*t*_207.59_ = −2.68, *P* = 0.008) and less fertile (*t*_211.25_ = −2.13, *P* = 0.034) eggs under thermal stress. In trials that progressed through the dispersal stage, larvae developed significantly faster in the harsh environment (6.04±1.47 days) compared to the benign (7.23±1.38; *t*_155.41_ = −5.24, *P* < 0.001) and were reduced for both number (F = 16.769, df = 1, 162, P < 0.001) and mass (F =22.346, df = 1, 156, P < 0.001; Fig 3). Parental breeding history had no effect on either larval number (F=4.458, df = 1, 162, P =0.409) or larval mass (F=0.019, df = 1, 156, P = 0.891).

**Figure 2:**
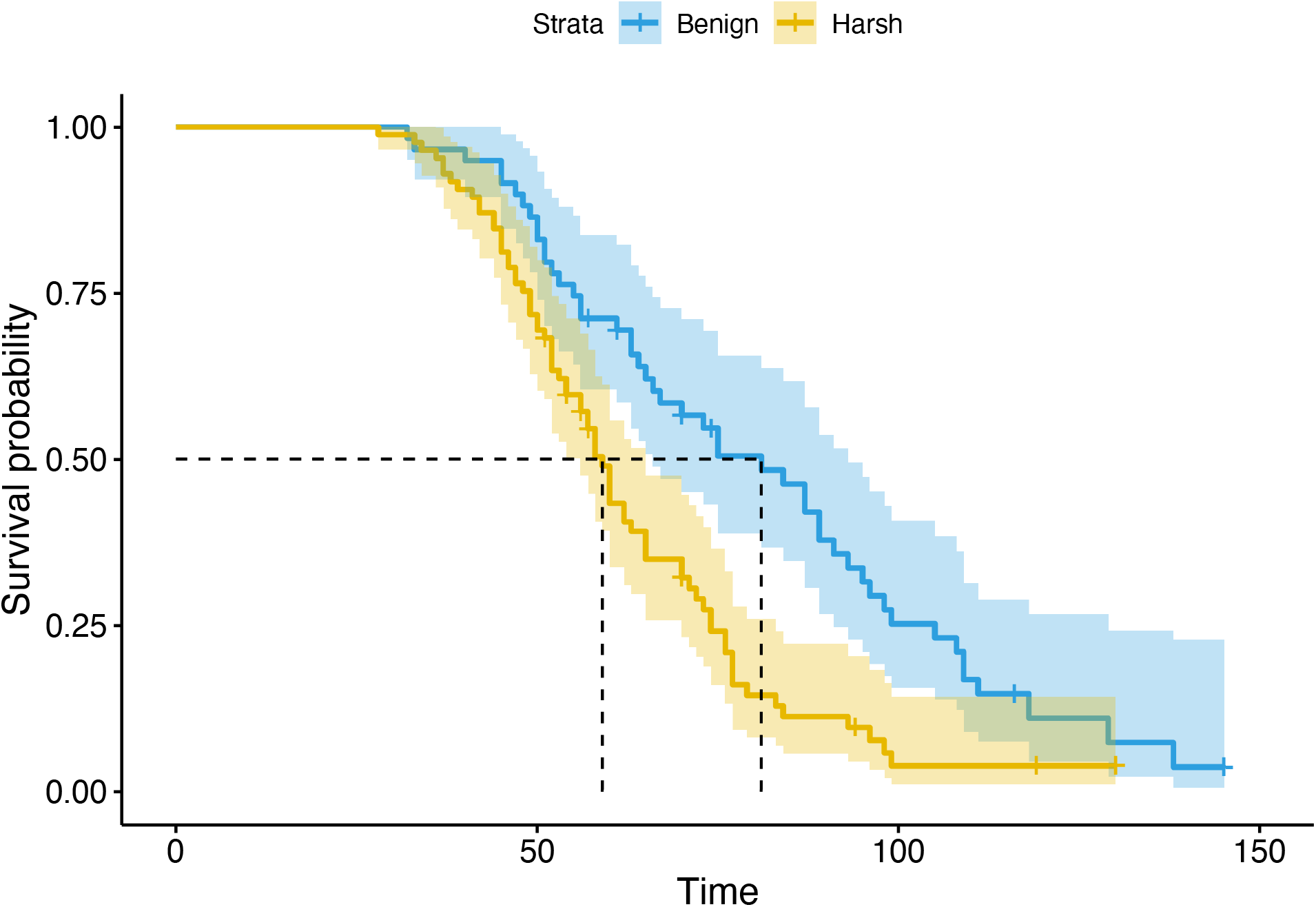
Survival curves calibrated from the mortality times of 144 beetles (25 censored) assigned to the benign (blue line; N = 59) and harsh (orange line; N = 85) thermal environments. Dotted lines indicate median lifespans for each environment, in days.

**Figure 3:**
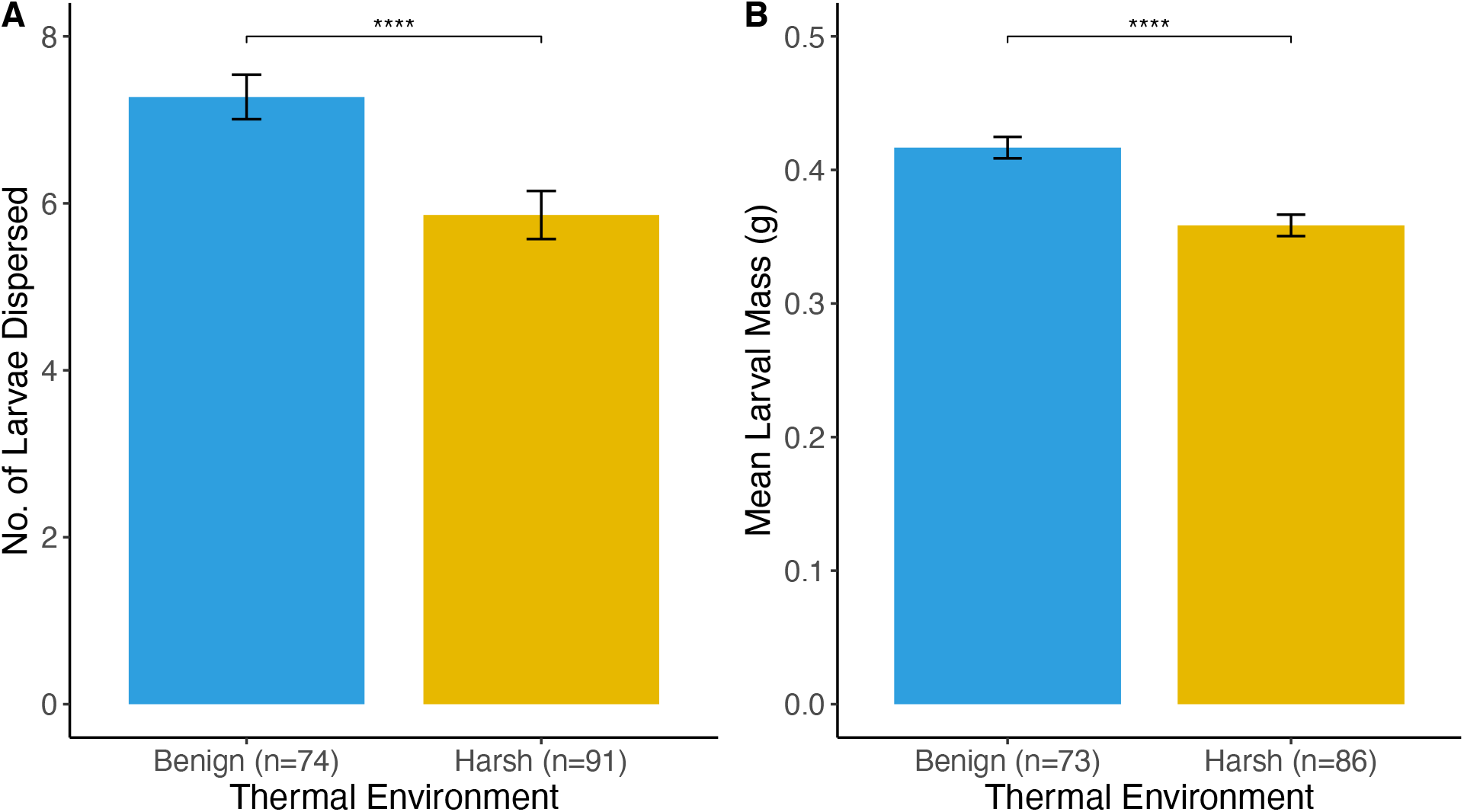
Comparisons of average breeding performance across all social conditions between the benign and harsh thermal environment. Performance is compared based on (A) number of larvae dispersed, and (B) mean larval mass (in g). Asterisks indicate significant differences at *α*=0.0001.

### Selection on Care

Visualizing selection, we found that care provided negligible fitness benefits under benign conditions (Fig 4A) and became harmful under stressful conditions (Fig 4B). Duration of parental attendance had significant linear effects on the number of offspring reared to dispersal in both environments (Table 1a, b). However, selection acting on distinct components of care differed between the two thermal environments. While there was no statistically significant selection acting on direct or indirect care in the benign environment, we found significant directional and stabilizing selection for different care components specific to the harsh environment. Total provisioning effort showed a positive linear relationship with brood size, whereas significant nonlinear effects on both larval size and number were detected in association with extreme values of indirect care. Thus, the harsh environment selected for increased time in direct provisioning and intermediate time in carcass maintenance. Variation was largely attributed to the number and sex of parents, as all behaviors varied significantly with social condition under both environmental conditions (Table 2a,b) and pairwise comparisons identified only indirect care as being significantly increased for biparental pairs relative to uniparental females (Table 3).

**Table 1:** Linear (*β*) and nonlinear (*γ*) standardized selection gradients (Lande and Arnold 1983, Brodie et al. 1995) relating brood performance (measured as number of larvae dispersed and mean larval mass (in g)) to behavioral measures of parenting effort. Selection gradients are presented separately for the (a) benign, and (b) harsh environment. Statistically significant gradients (at α=0.05) are highlighted in bold.

| Behavioral Measure | N | Larval Number |  |  |  | Mean Larval Mass |  |  |  |  |
| --- | --- | --- | --- | --- | --- | --- | --- | --- | --- | --- |
| | | $\beta$ | $P$ | $\gamma$ | $P$ | N | $\beta$ | $P$ | $\gamma$ | $P$ |
| **(a) Benign Environment** |  |  |  |  |  |  |  |  |  |  |
| Days Attended | 74 | **0.124**<br>($\pm 0.034$) | <0.001 | -0.259<br>($\pm 0.078$) | 0.078 | 73 | 0.028<br>($\pm 0.021$) | 0.179 | -0.006<br>($\pm 0.050$) | 0.907 |
| Total Indirect Care | 74 | 0.058<br>($\pm 0.035$) | 0.097 | -0.025<br>($\pm 0.052$) | 0.632 | 73 | -0.004<br>($\pm 0.020$) | 0.832 | -0.005<br>($\pm 0.032$) | 0.867 |
| Total Direct Care | 74 | 0.042<br>($\pm 0.034$) | 0.228 | 0.008<br>($\pm 0.050$) | 0.868 | 73 | 0.028<br>($\pm 0.019$) | 0.164 | -0.021<br>($\pm 0.031$) | 0.513 |
| **(b) Harsh Environment** |  |  |  |  |  |  |  |  |  |  |
| Days Attended | 91 | **0.161**<br>($\pm 0.052$) | 0.002 | -0.089<br>($\pm 0.131$) | 0.497 | 86 | 0.039<br>($\pm 0.027$) | 0.167 | 0.035<br>($\pm 0.068$) | 0.610 |
| Total Indirect Care | 90 | -0.001<br>($\pm 0.052$) | 0.992 | **-0.165**<br>($\pm 0.069$) | 0.020 | 85 | -0.037<br>($\pm 0.027$) | 0.179 | **-0.100**<br>($\pm 0.035$) | 0.006 |
| Total Direct Care | 90 | **0.102**<br>($\pm 0.051$) | 0.049 | 0.008<br>($\pm 0.060$) | 0.896 | 85 | 0.021<br>($\pm 0.026$) | 0.416 | -0.053<br>($\pm 0.029$) | 0.069 |

**Table 2:** Mixed model ANOVAs testing the effects of social condition (uniparental male, uniparental female, and biparental) on parental effort (the total time allocated to indirect or direct care during a 30-minute observation) for three forms of care. Results are reported separately for the benign and harsh environment, with male and female IDs treated as random factors.

| Model | MS | Num $df$ | Den $df$ | $F$ | $P$ |
| --- | --- | --- | --- | --- | --- |
| **(a) Benign Environment** |  |  |  |  |  |
| Days Attended | 20.06 | 2 | 71.00 | **8.60** | < 0.001 |
| Total Indirect Care | 5.00 | 2 | 37.75 | **12.34** | < 0.001 |
| Total Direct Care | 96.68 | 2 | 40.05 | **6.22** | 0.004 |
| **(b) Harsh Environment** |  |  |  |  |  |
| Days Attended | 14.37 | 2 | 57.81 | **15.65** | < 0.001 |
| Total Indirect Care | 18.05 | 2 | 61.10 | **27.36** | < 0.001 |
| Total Direct Care | 124.98 | 2 | 63.29 | **3.20** | 0.047 |

**Table 3:** *a priori* determined pairwise comparisons of parental effort among pairs within the same acclimation environment, benign or harsh. For each dataset, biparental (BP) is the reference group against which uniparental female (UPF) and uniparental male (UPM) observations are contrasted. Effects with statistically significant *P-*values (at α=0.05) are shown in bold.

| *Contrast* |  | *t* | *P* |
| --- | --- | --- | --- |
| ***Days Attended:*** |  |  |  |
| Benign Environment | BP – UPF | -0.836 | 0.684 |
|  | **BP – UPM** | 3.080 | 0.010 |
| Harsh Environment | BP – UPF | 0.069 | 0.997 |
|  | **BP – UPM** | 4.967 | < 0.001 |
| ***Total Indirect Care:*** |  |  |  |
| Benign Environment | **BP – UPF** | 3.885 | 0.001 |
|  | **BP – UPM** | 4.119 | < 0.001 |
| Harsh Environment | **BP – UPF** | 3.952 | 0.006 |
|  | **BP – UPM** | 7.127 | < 0.001 |
| ***Total Direct Care:*** |  |  |  |
| Benign Environment | BP – UPF | 1.250 | 0.430 |
|  | **BP – UPM** | 3.408 | 0.005 |
| Harsh Environment | BP – UPF | 0.391 | 0.919 |
|  | BP – UPM | 2.328 | 0.064 |

**Figure 4:**
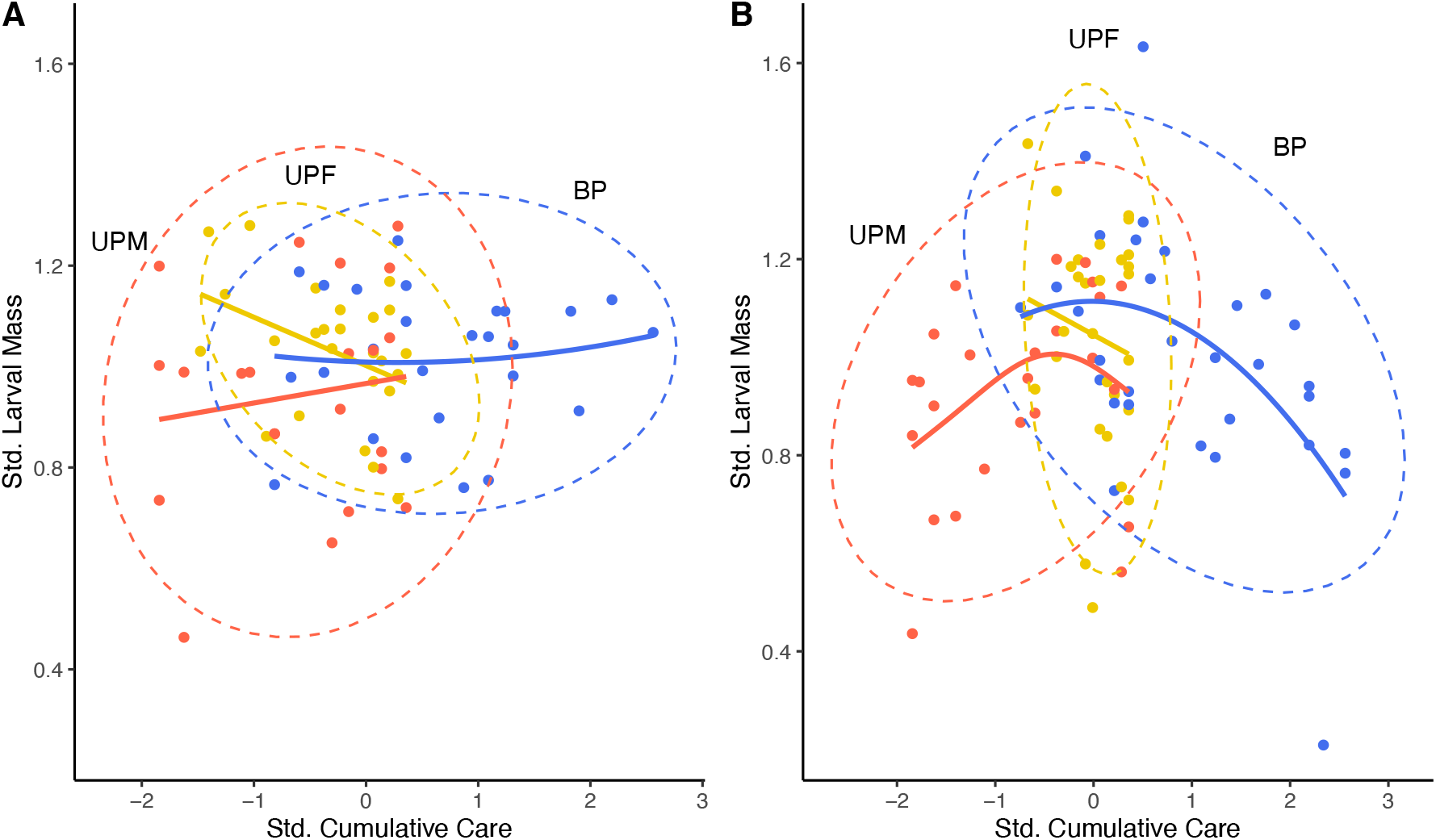
Relationship between standardized total parental time allocation to care (direct + indirect) and standardized mean larval mass in the (A) benign environment, and (B) harsh environment. Data points belonging to each of three social conditions (uniparental male (UPM), uniparental female (UPF), and biparental (BP)) are differentiated by color (red = UPM, yellow = UPF, blue = BP). Labeled ellipses illustrate 95% confidence intervals, and approximate splines illustrate overall trends in the data.

### Plasticity of Biparental Care

For plasticity of biparental care to help mitigate increased environmental costs of parenting, males and females should adjust investment strategies to complement, rather than overlap, with care components of their partner. We found sex-specific tradeoffs in parenting behaviors, consistent with covariance underpinning potential responses. Repeated measures ANOVA tests identified social condition as an important factor explaining within-subject variation of parenting behavior, but these effects were sex-specific (Table 3). Females were not plastic and appeared to care at capacity regardless of thermal environment or social condition. Conversely, male parenting effort varied significantly between social conditions, most dramatically in terms of duration of care (Table 4). Further, males in the harsh environment significantly reduced their provisioning effort when paired with a female but maintained the same levels of indirect care (Table 4). Between-treatment comparisons largely recapitulated these sex-specific patterns: social condition emerged as a significant predictor of overall within-subject behavioral variation in males (WTS = 15.691, *P* < 0.001) but not in females (WTS = 4.816, *P* = 0.185). Neither males nor females showed significant differences in parenting behavior between thermal environments (Males: WTS = 0.399, *P* = 0.945; Females: WTS = 3.752, *P* = 0.295). Moreover, adaptive plasticity of biparental care did not mitigate effects of environmental stress because the interaction of within-treatment (social condition) and between-treatment (thermal environment) effects was not significant.

**Table 4:** Repeated-measures ANOVAs quantifying behavioral plasticity of individual female and male parenting behaviors (days in attendance and time allocation to indirect and direct care during a 30-minute observation). Test statistics are reported separately for each sex and thermal environment (benign and harsh), where social condition (uniparental or biparental) is the within-subject factor and individual ID nested within trial number is the error term. Effects with statistically significant *P-*values (at α=0.05) are shown in bold.

| Behavior | Test statistic | *Benign* |  | *Harsh* |  |
| --- | --- | --- | --- | --- | --- |
|  |  | Female | Male | Female | Male |
| Days Attending | *F* (df) | 0.730 (1,19) | **18.4 (1,19)** | 3.307 (1,19) | **4.994 (1,17)** |
|  | *P* | 0.403 | < 0.001 | 0.085 | 0.039 |
| Indirect Care | *F* (df) | 0.231 (1,19) | 0.699 (1,19) | 0.632 (1,19) | 0.252 (1,17) |
|  | *P* | 0.637 | 0.413 | 0.436 | 0.622 |
| Direct Care | *F* (df) | 0.101 (1,19) | 3.165 (1,19) | 0.012 (1,19) | **4.754 (1,17)** |
|  | *P* | 0.755 | 0.091 | 0.912 | 0.044 |

## Discussion

In this study, we investigated the potential of plasticity of biparental care to ameliorate a harsh environment in a burying beetle, *Nicrophorus orbicollis*. Our prediction was that offspring receiving more care through additive or load lightening benefits of multiple caregivers would fare better under harsh environmental conditions. We tested this by exposing families with different parental compositions to thermal stress and identifying behavioral correlates of performance using standardized selection gradients. The patterns that emerged were opposite to expectations: the harsh environment favored intermediate, not high overall parental investment, but the type of care was important, and components were not independent of each other. Plasticity of biparental care did not help overcome constraints because decision rules for investment were sex-specific and were unaltered by generalized stress on the family. The consequence was that in biparental pairs under environmental stress, females overinvested, and males contributed only to care types that caused a decrease in fitness in excess. These results challenge our understanding of the adaptive role of biparental care in hostile environments.

The thermal stress we imposed had strong deleterious fitness effects compared to a more benign temperature. Not only did adults acclimated to the warmer environment suffer reduced lifespans and lower reproductive potential per bout, but offspring were also less likely to survive to dispersal and attained lower body mass than counterparts in the benign environment. Our results are consistent with both field and laboratory studies of the genus noting significant performance declines along gradients of temperatures (Meierhofer et al. 1999, Müller et al. 2007, Steiger et al. 2007, Jacques et al. 2009, Quinby 2016, Ong 2019, Feldman 2020). Given these severe fitness costs, we expected there would be selection pressure to cope with extreme temperatures.

Because environmental hostility exacerbates offspring vulnerabilities, systems with dependent young should benefit from the capacity to increase parental investment when confronted with more extreme environments (Wilson 1975, Wesolowski 1994, 2004). Burying beetles are known for the level of plasticity they show in parental care, especially in response to social condition, with uniparental female, uniparental male, and biparental care possible for many species including *N. orbicollis* (Trumbo 1991, Scott 1998a, Smiseth and Moore 2004, Suzuki and Nagano 2009, Benowitz et al. 2016). We predicted that given this extant plasticity, care by two parents would have greater capacity to respond to increased offspring need. Contrary to this expectation, our high-stress environment did not induce strong and consistent directional selection relative to the benign environment. Instead, overall care was associated with significant nonlinear effects – an indication of strong stabilizing selection (Schluter 1988). This translated to fewer and smaller offspring among caregivers with both the lowest and the highest cumulative behavioral investments (Fig 4B). We detected no improvements in performance among families with two caregivers as opposed to one (Fig 4B). In fact, because two caregivers are effectively capable of twice the sum total effort, biparental pairs accounted for much of the performance reduction in the upper tails of the care distribution. These results are consistent with independent investigations carried out in Oregon (Feldman 2020) and Canada (Ong 2019), which report significantly reduced performance and limited compensation among biparental pairs exposed to experimental warming treatments. Our study provides a mechanism for these effects: reduced offspring performance at higher temperatures does not result from biparental care *per se,* but from temperature-dependent thresholds in optimal care allocation, which are most likely to be exceeded when two parents are active at the nest.

Our original prediction – that parents should compensate for environmental stress by increasing the amount of time they spend caring – was based on an assumption of unconstrained potential to evolve more care to offspring. Clearly parents should have the ability to re-allocate time from non-caring to caring behaviors, as males can shift to care from no-care when females are not present. However, care in burying beetles comes in different forms, and our analysis suggests that there is conflicting selection resulting in a constraint on changes in care composition. In the harsh environment, increased direct care was under directional selection, with more care associated with greater offspring number and mass. In contrast, indirect care was under stabilizing selection for both offspring number and mass (Table 1). It is possible that negative genetic correlations between indirect care and direct care, as described in quantitative genetic work in the related *N. vespilloides* (Walling et al. 2008), limit independent responses of care components under conflicting selection regimes. However, such tradeoffs would not explain the lack of adaptive plasticity seen in biparental pairs.

Our expectation was that because burying beetles show flexibility in parenting in response to social parameters (Trumbo 1991, Smiseth et al. 2005, Suzuki and Nagano 2009, Creighton et al. 2015, Mattey and Smiseth 2015, Parker et al. 2015), and males caring with females have spare capacity, the application of a generalized stressor should promote shifts from conflict to cooperation. Instead, we found that both males and females adhered to predicted sex-specific rules for parental investment – males were plastic and females were not, as seen in the related *N. vespilloides* (Smiseth et al. 2005, Royle et al. 2014) – irrespective of the selective environment. Specifically, females cared at capacity even when exposed to heat stress and provided with male helpers, allowing us to reject any ‘load lightening’ benefits of two caregivers (Crick 1992, Johnstone 2011). Males in the presence of females withheld direct care and deserted broods earlier, consistent with sexual conflict. The main difference between biparental pairs and uniparental females across environments was an increase in indirect care, underpinned by the fact that neither males nor females showed plasticity of this behavior. Reductions in offspring fitness with additive care were most likely due to tradeoffs with direct care at the individual level. Hence, when ambient conditions turn unfavorable, two parents are not more efficient at caring for offspring than one.

Given the predominance of biparental care in *N. orbicollis*, why have mechanisms for enhancing coordination not evolved? In systems where brood care responsibilities are shared by more than one individual, social factors are expected to have an outsized influence on investment decisions, and the ability to mount coordinated responses may help buffer environmental variation (Heinsohn 2004, Ridley and Raihani 2008). Models of biparental care such as partial compensation (Houston and Davies 1985), negotiation (McNamara et al. 1999), and turn-taking (Johnstone et al. 2014) assume that male and female strategies are optimized to resolve conflict over offspring care. However, if biparental care of burying beetles did not evolve to mitigate offspring need, then the dynamics predicted under these models may not hold true. As in the related *N. vespilloides*, sexual conflict may continue to structure interactions between the sexes (Boncoraglio and Kilner 2012, Parker et al. 2015), at least in terms of a lack of a response by males to offspring need. In burying beetles, the prevailing theory for why males of some species join females is that mating opportunities are limited elsewhere and pairs fare better than single parents in defense of the valuable but temporally and spatially unpredictable resources (Scott 1990, 1994, 1998a, Trumbo 1991). If true, then the impetus for the evolution of care may be different for males and females. While females should focus efforts on meeting the needs of offspring, males should remain impervious to offspring needs except in the extreme case that the female dies or abandons the nest. Thus, males adopt an ‘insurance policy’ strategy for participation of care (Parker et al. 2015). This is supported by evidence in a variety of burying beetle species for compensatory responses to partner removal (Smiseth et al. 2005, Suzuki and Nagano 2009, Royle et al. 2014) and no compensation for subtler perturbations in the family environment, such as reduced partner provisioning, (Suzuki and Nagano 2009, Suzuki 2020), increased offspring begging (Suzuki 2020) or, as we show here, the application of a generalized stressor. Overall, research on burying beetles suggest that there can be sex-specific evolutionary pathways for biparental care consistent with sexual conflict as one of the drivers of the evolution of care in this genus (Boncoraglio and Kilner 2012, Parker et al. 2015).

The prediction that cooperative parental strategies enhance resilience in harsh or hostile environments is not novel (Wilson 1975, Emlen 1982), but climate change has afforded new urgency to understanding its practical significance (Lucey et al. 2015, Manfredini et al. 2019, Henriques and Osmond 2020). Our study has shown that burying beetles at southern range margins will face steep reproductive challenges associated with rising temperatures alone, and that these will not be alleviated through biparental cooperation. Despite the predominance of biparental social structures in this species strategies for coordinated care are unrefined. The implication of our work is that the potential for parenting to ameliorate the effects of climate change is likely to depend on the evolutionary drivers of parental care, which may be sex specific and be best reflected in components of care.

## Acknowledgements

K.E. Kollars and the beetle crew assisted with colony maintenance. We thank C.B. Cunningham, P.J. Moore, E.A. Shelby, S.D. Harris, E.A. McKinney, and J.T. Washington for constructive comments and feedback. J.B.M. was supported by a USDA-ARS cooperative agreement (#58-6080-9-006).

## Notes

**Conflict of Interest Statement:** The authors declare no conflicts of interest.

### Competing Interest Statement

The authors have declared no competing interest.

