## Supplemental Figure for "Constrained flexibility of parental cooperation limits evolutionary responses to harsh conditions"

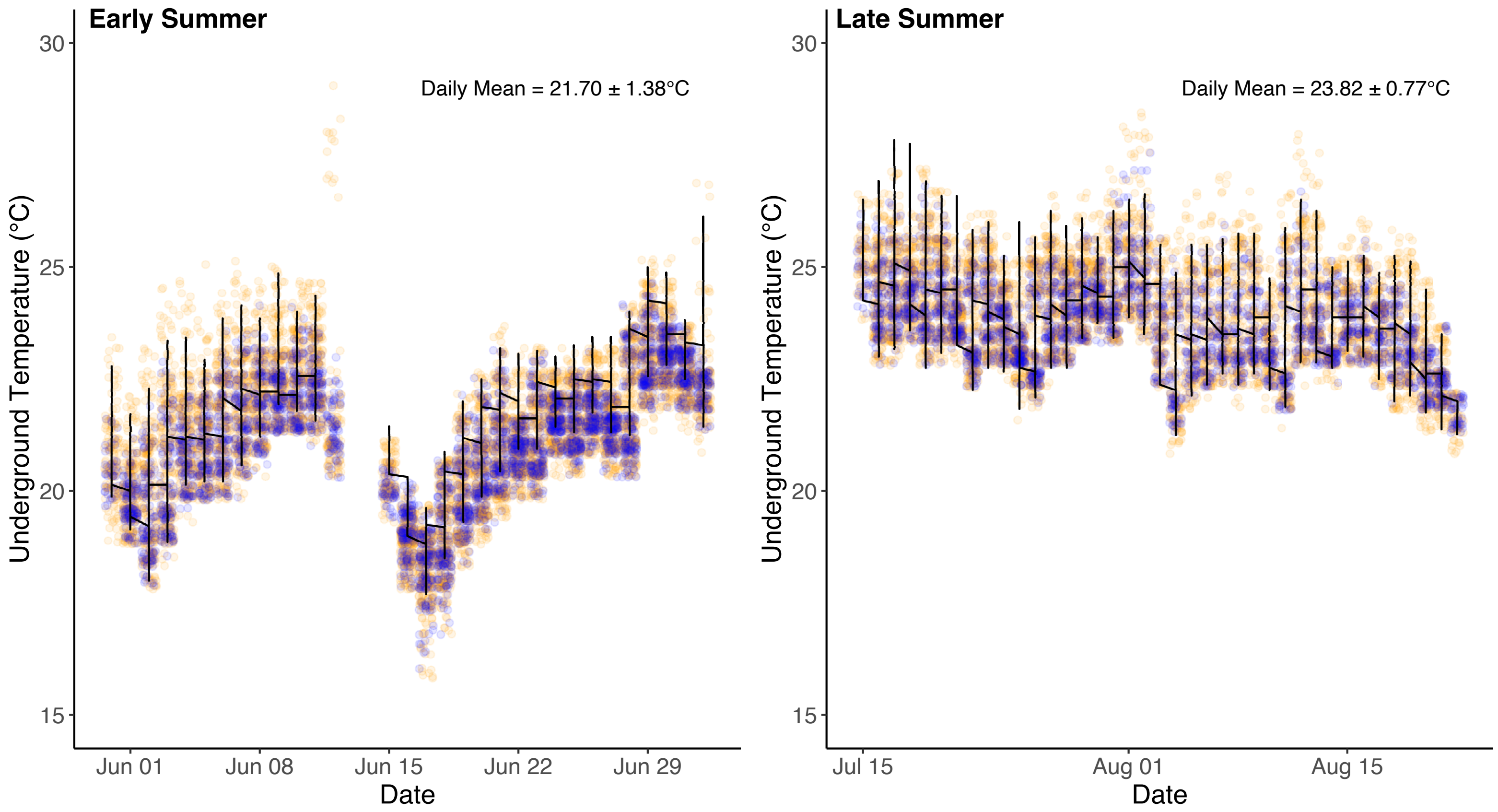


Figure S1: Trends in ambient temperature in Whitehall Forest, Athens, GA measured over two periods of the active season of *Nicrophorus orbicollis*: between 31 May–03 July 2020 ('Early Summer'; top) and 15 July–22 August ('Late Summer'; bottom). Temperature data were collected every 30 minutes using Thermochron® iButton temperature loggers (©Maxim Integrated Products, Inc., San Jose, CA, U.S.A) deployed 10-12 cm underground at beetle trap locations. This was done to ensure that the microenvironments where burying beetles are most likely to breed (underground, beneath the forest canopy) were represented. In each panel, points represent temperature recordings, where daytime temperatures (0700-2000) are shown in orange and nighttime temperatures (2000-0700) are shown in blue. Lines represent trends in daily mean temperature.
